# Viral antigen mismatch affects antiviral T-cell response and may impair immunotherapeutic efficacy against ATL

**DOI:** 10.1101/2024.01.25.576615

**Authors:** Kenji Sugata, Mitsuyoshi Takatori, Omnia Reda, Benjy Jek Yang Tan, Masahito Tokunaga, Tomoo Sato, Mitsuharu Ueda, Yoshihisa Yamano, Atae Utsunomiya, Yorifumi Satou

## Abstract

Human T-cell leukemia virus type 1 (HTLV-1) has the potential to transform primary CD4^+^ T cells *in vitro* within a short time; however, the majority of infected individuals maintain an asymptomatic and disease-free condition, suggesting the existence of an equilibrium between the proliferation of infected cells and host immunity. The decline in anti-viral immunity contributes to the transformation of the infected cells, leading to the development of adult T-cell leukemia/lymphoma (ATL). This study identified a variation in a major viral antigen, HTLV-1 Tax, in human leukocyte antigen-A24 (HLA-A24) positive individuals. Two variants of Tax_301-309_ peptides, SFHNLHLLF (Tax_301-309_ A) and SFHSLHLLF (Tax_301-309_ B) were found to induce distinct T-cell immune responses in HLA-A24 positive individuals. There was a disparity between two Tax_301-309_ peptides in the detection of anti-Tax_301-309_ cytotoxic T-lymphocytes (CTLs) binding to A24/peptide multimers by flow cytometry analysis. More importantly, over half of the anti-Tax TCRs of anti-Tax CTLs from infected individuals did not recognize mismatched Tax_301-309_ peptides by Enzyme-Linked Immunospot (ELISpot) assay using Jurkat T cells expressing the anti-Tax_301-309_ specific TCR. These findings underscore the importance of matching the viral antigen epitope type in T-cell-based immunotherapy against ATL by using viral antigen Tax.

**Key points:**

- Epitope heterogeneity in the major viral antigen in HTLV-1 infection causes different T-cell responses in infected individuals.
- Recommended guideline; performing virus typing to obtain optimal efficacy in T-cell-mediated immunotherapy against the viral antigen Tax

## Introduction

Human T-cell leukemia virus type 1 (HTLV-1) is a human retrovirus, which induces chronic and persistent infection in humans. In stark contrast with another human retrovirus HIV-1, viral particles are rarely detected in the plasma of HTLV-1 infected subjects; however, there are 1 – 2 % of infected cells circulating in the peripheral blood of these individuals^1^. Both cellular and humoral anti-viral immune responses are detectable in most infected individuals, indicating minimal but persistent viral gene expression. Considering the potent transformation capacity of HTLV-1 *in vitro*^*2*^, there should be an equilibrium between the proliferation of infected cells and host immunity. This equilibrium maintains an asymptomatic phase in most of the infected individuals, preventing the onset of transformation of the infected cells and the development of adult T-cell leukemia/lymphoma (ATL). This notion is supported by several pieces of clinical evidence such as the high frequency of ATL onset in immunocompromised host^3,4^. Additionally, there is a high frequency of anti-viral CTLs detected in ATL patients with a good clinical course after stem cell transplantation^5,6^. As a therapeutic approach for ATL, T-cell-based immunotherapy against ATL by using viral antigen Tax is currently under preclinical and clinical development ^7^.

There are several strains of HTLV-1 reported^8^, but the sequence of HTLV-1 is generally conserved when compared with that of HIV-1. This is because HIV-1 proliferates mainly via viral replication whereas HTLV-1 replicates through cell mitotic proliferation. However, we recently identified sequence heterogeneity in immune-dominant HTLV-1 protein Tax^9,10^. This heterogeneity may affect T-cell immunity against the virus, raising an important caution for immunotherapy in HTLV-1-induced leukemia/lymphoma.

## Study design

### Ethics statement

All protocols involving human subjects were reviewed and approved by the Kumamoto University Institutional Review Board (approval number 263). This study was carried out in accordance with the guidelines proposed in the Declaration of Helsinki. Informed written consent was obtained from all subjects in this study.

### Cell culture and reagents

Tax_301-309_-A (SFHNLHLLF) and -B (SFHSLHLLF), Nef_134-143_ (RYPLTFGWCF) and NY-ESO1_157–165_ (SLLMWITQV) were purchased from Genscript and dissolved at 50 mg/ml in DMSO. PBMCs were isolated from whole blood using Ficoll-Paque (GE Healthcare Life Sciences).

### Tax-A and -B genotype in HTLV-1 genome

We evaluated the differences of provirus sequences among individuals by aligning previously performed HTLV-1 DNA-capture-seq data (98 HTLV-1-infected individuals) to a reference sequence (GenBank: AB513134)^11^.

### HLA reconstitution assay

Peptide reconstitution assay was performed as shown previously^12^. Briefly, HLA-A24-transduced K562 cells were treated with an acid solution and then incubated in the presence of β2-microglobulin and peptide. HLA reconstitution was evaluated by staining with anti-HLA-A24 monoclonal antibody and using flowcytometry. The EC_50_ was defined as the concentration of peptide required to achieve 50% of the maximal response.

### HLA-A24/Tax multimer staining

HLA-A24/Tax tetramer and dextramer were obtained from MBL and Immudex Co. Ltd, respectively. Whole PBMCs were first stained with HLA-A24/Tax tetramer and dextramer. Following multimer staining, the cells were stained by anti-CD8 monoclonal antibody and LIVE/DEAD Fixable Near-IR Cell Stain Kit. The stained cells were analyzed with flow cytometry.

### TCR transduction

The TCR sequence of HT-9 cells has been reported in a previous study^13^. We performed single cell TCR-seq analysis with 10x Chromium system to acquire TCR sequence information of anti-Tax_301-309_ CTL according to our previous report with minor modifications^14^.

To identify anti-Tax_301-309_ CTL in the single cell analysis, we utilized a custom-made Tax-peptide-HLA dextramer with 10x barcode. We obtained 32 TCRαβ sequences from individuals infected with either Tax type A or Tax type B HTLV-1. The TCRαβ pairs were fused with P2A (self-cleaving 2A peptides) and retrovirally transduced into TCR-deficient Jurkat cells^15^.

### ELIS_POT_ assay

TCR-transduced Jurkat cells were stimulated by HLA-A24/K562 cells pulsed with peptide at 10 μg/ml. Specific cytokine production was assessed using IL-2 ELIS_POT_ Kit (Mabtech) and the ELIPHOTO Counter (Minerva Tech). The EC_50_ was defined as the concentration of peptide required to achieve 50% of the maximal response.

### Statistical analysis

Data were analyzed using a chi-squared test or Unpaired two-tailed Student’s t-test with GraphPad Prism 7 software (GraphPad Software Inc.) unless otherwise stated. Statistical significance was defined as P < 0.05.

### Data Sharing Statement

Single cell TCR-seq data will be deposited to public database before publication.

## Results and Discussion

In a previous study, we determined the whole HTLV-1 sequence using DNA-capture methods^11^. Our analysis revealed two types of Tax sequences with differing pathogenicity, consistent with previous report by others (Figure 1A). Additionally, a nucleotide variation in the coding region for Tax was identified in our previous study^16^ (Figure 1B). Importantly, the nucleotide variation is associated with an amino acid difference in the anti-viral cytotoxic T-lymphocyte (CTL) epitope HTLV-1 Tax_301-309_. This epitope is a major viral CTL epitope for HLA-A*24:02 (A24)^17^, the most frequent HLA-A type in Japan and some other HTLV-1 endemic areas (Figure 1C). The two types of Tax_301-309_ peptides are SFHNLHLLF (Tax-A_301-309_) and SFHSLHLLF (Tax-B_301-309_). Notably, the variable amino acid position in the epitope is not responsible for binding to HLA^18^. We also experimentally confirmed that there was no significant difference in binding activity to HLA-A24 between the two Tax_301-309_ variants using HLA reconstitution assay (Figure 1D). To analyze antigen specific CTLs, flow cytometry was performed using A24/ Tax_301-309_ multimers. Despite the availability of several A24/Tax_301-309_ multimers commercially, it’s worth noting that they only contain Tax-B_301-309_. Therefore, we ordered custom-made A24/Tax_301-309_ multimers containing Tax-A_301-309_ and analyzed their detection efficiency for anti-Tax_301-309_ CTL. Flow cytometry analysis of PBMCs from three individuals infected with either Tax-A_301-309_ or Tax-B_301-309_ type HTLV-1 revealed that virus-type matched tetramer assays showed a higher frequency of tetramer-positive CD8^+^ cells with high mean fluorescence intensity (MFI) value whereas mismatched ones showed lower frequency of tetramer-positive CD8^+^ cells with low MFI value (Figure 2A). Similar result was observed with A24/Tax_301-309_ dextramer staining (Figure 2B). These data indicate that virus-type matched A24/Tax_301-309_ multimers have stronger binding affinity to TCR of anti-Tax_301-309_ CTL than mismatched ones. Using mismatched A24/Tax_301-309_ multimers could potentially lead to an underestimation of the frequency of anti-Tax CTL in the assay. Next, we performed single cell TCR-seq with 10x Chromium system to obtain TCR sequence information of anti-Tax_301-309_ CTL. We ordered custom-made A24/ Tax-A/B_301-309_ dextramer containing 10x barcode to identify anti-Tax_301-309_ CTL in the single cell analysis. Thirty-two TCR sequences of anti-Tax_301-309_ CTL were obtained from HTLV-1 individuals infected with either Tax-A or Tax-B type of HTLV-1. We used Jurkat T cells without expressing TCR on the cell surface to generate specific T cells expressing anti-Tax_301-309_ TCR. Subsequently, we performed Enzyme-Linked ImmunoSpot (ELISpot) assay to evaluate each CTL’s ability to produce cytokine upon encountering a specific antigen. Consistent with A24/Tax multimer assay, Jurkat T cells with TCR from Tax-A infected individuals showed higher activity against SFHNLHLLF (Tax-A_301-309_) than SFHSLHLLF (Tax-B_301-309_) (Figure 2C). In contrast, Jurkat T cells with TCR from Tax-B infected individuals showed higher activity against Tax-B_301-309_ than Tax-A_301-309_. Among the 32 anti-Tax_301-309_ TCRs, more than half, 56.2%, were able to respond exclusively to the virus-type matched peptide (Figure 2D). One-fifth of them exhibited partial responsiveness, while others were equally responsive to mismatched Tax_301-309_ peptide (Figure 2D), consistent with the finding that there was no significant difference between A24/Tax A_301-309_ and A24/Tax B_301-309_ multimer-positive cells in some infected individuals (Figure 2A and B). We further investigated the binding avidity of anti-Tax_301-309_ TCRs to Tax-A/B_301-309_ by cultivating them with different concentrations of peptides with antigen presenting A24/K562 cells (Figure 3A). We analyzed two anti-Tax_301-309_ specific TCRs (Tax-TCR-1 and -2) with cross-reactivity between two types of Tax_301-309_ and HT-9 TCR from a previous report by others^13^. HT-9 showed strong binding avidity to Tax_301-309_ peptide with Tax A type but decreased binding to Tax_301-309_ peptide with Tax B (Figure 3B). These data indicate that using mismatched TCRs for T-cell therapy and Tax peptides for ATL treatment may not have enough effect due to antigen peptide mismatch.

**Figures 1.**
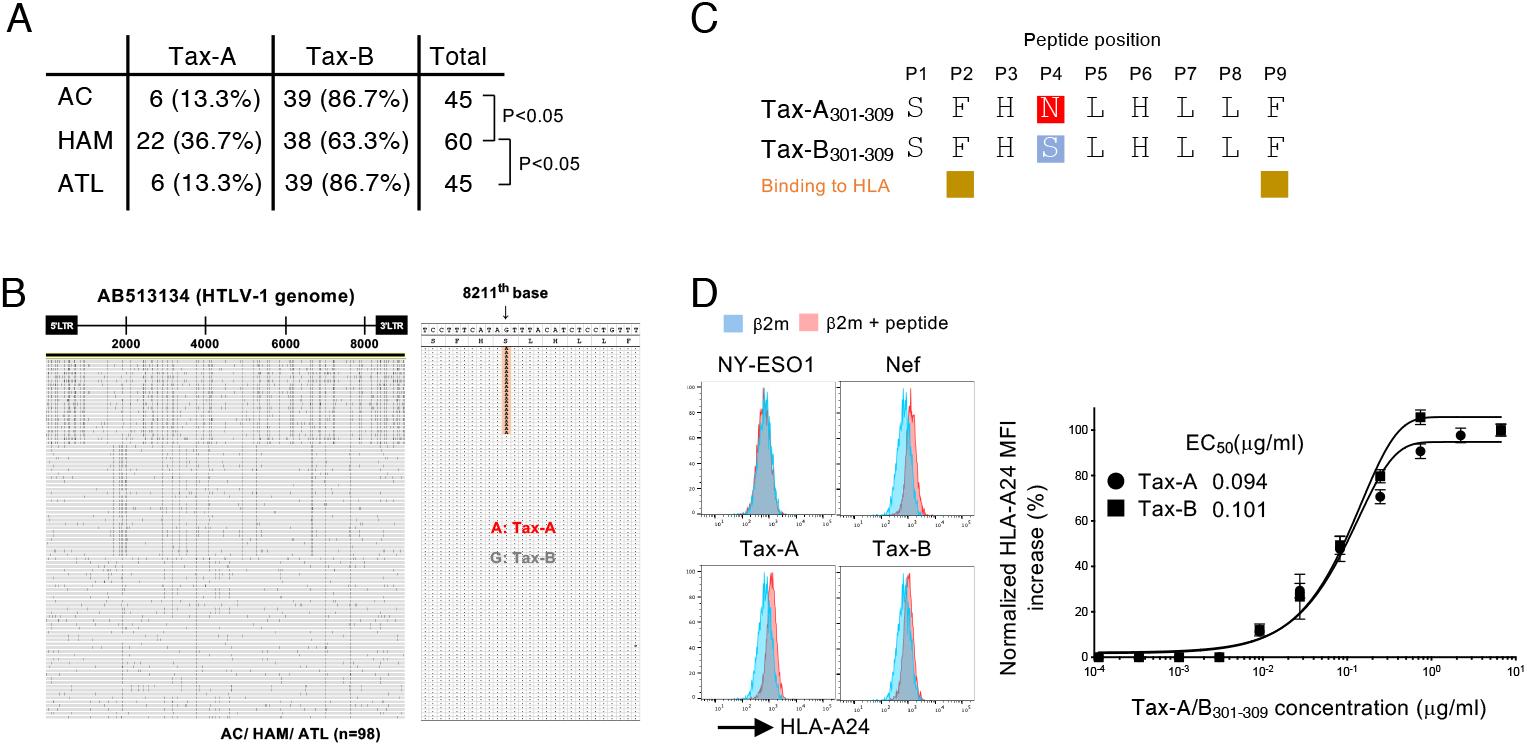
Asparagine and Serin substitution at position 304 of HLA-A*24:02-restricted Tax_301-309_. (A) List for percentages of Tax-A and -B in HTLV-1 infected patients. (B) Differences of HTLV-1 provirus sequences among individuals. The provirus sequences of 98 HTLV-1-infected cases obtained through HTLV-1 DNA-capture-seq were aligned to the reference sequence (AB513134). Nucleotides differing from the reference sequence are highlighted in black. Nucleotide substitutions were detected at the 8211^th^, where A corresponds to Tax-A and G corresponds to Tax-B. (C) Comparison of amino acid sequences between Tax-A and -B with HLA-A*24:02 restriction. Note that anchor residues for HLA-A24 binding in the peptides are P2 and P9. (D) Tax peptide reconstitution assay for HLA-A24 restriction. Acid buffer-treated HLA-A24/K562 cells were incubated with peptide and beta-2-microglobulin. HLA-A24/peptide reconstitution was evaluated by using anti-HLA-A24 antibody and flow cytometry. NY-ESO1_157-165_ and Nef_134-143_ were negative and positive control as HLA restriction, respectively. The EC_50_ was defined as the concentration of peptide required to achieve 50% of the maximal response. Dots and error bars represent the mean ± SD of results in triplicate experiments, and at least 2 independent experiments were performed.

**Figures 2.**
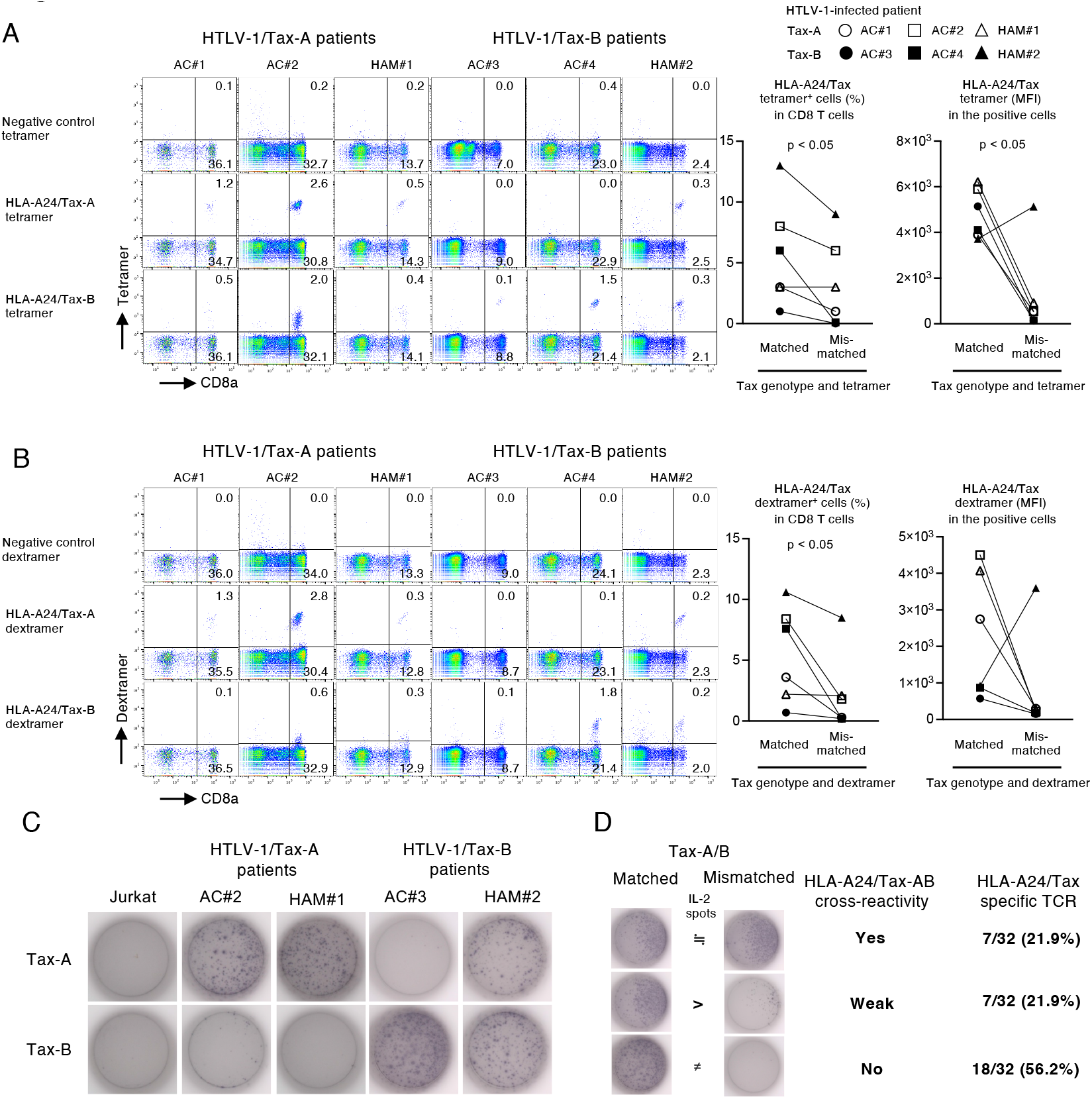
Most of specific TCRs recognize either Tax-A or -B peptides with HLA-A*24:02 restriction. (A and B) HLA multimer staining for HLA-A24/Tax specific TCRs in CD8 T cells. HLA-A24^+^ six HTLV-1-infected patients (four ACs and two HAM) were grouped as Tax-A and -B based on HTLV-1 genotype. PBMCs from the patients were stained with tetramer (A left) and dextramer (B left) for Tax-A or -B peptides with HLA-A*24:02 restriction. The multimer positive cells (%) and the mean fluorescent intensity (MFI) for each patient were compared between matched and mis-matched tetramer (A right) and dextramer (B right) staining. (C) IL-2 ELIS_POT_ assay for HLA-A24/Tax specific TCR. Four TCR pairs were newly cloned from CD8 T cells in peripheral blood of ACs and HAM patients. TCR-deficient Jurkat cells were transduced with specific TCRs and stimulated by HLA-A24/K562 cells pulsed with Tax-A and -B peptides. (D) Summary of TCR cross-reactivity for HLA-A24/Tax-A and -B. Thirty-two specific TCRs newly cloned from ACs and HAM patients were stimulated with peptide-pulsed HLA-A24/K562 cells for IL-2 ELIS_POT_ assay. The TCR cross-reactivity was evaluated by comparing the number of IL-2 spots in TCR-transduced Jurkat cells after peptide stimulation. At least 2 independent experiments were performed.

**Figures 3.**
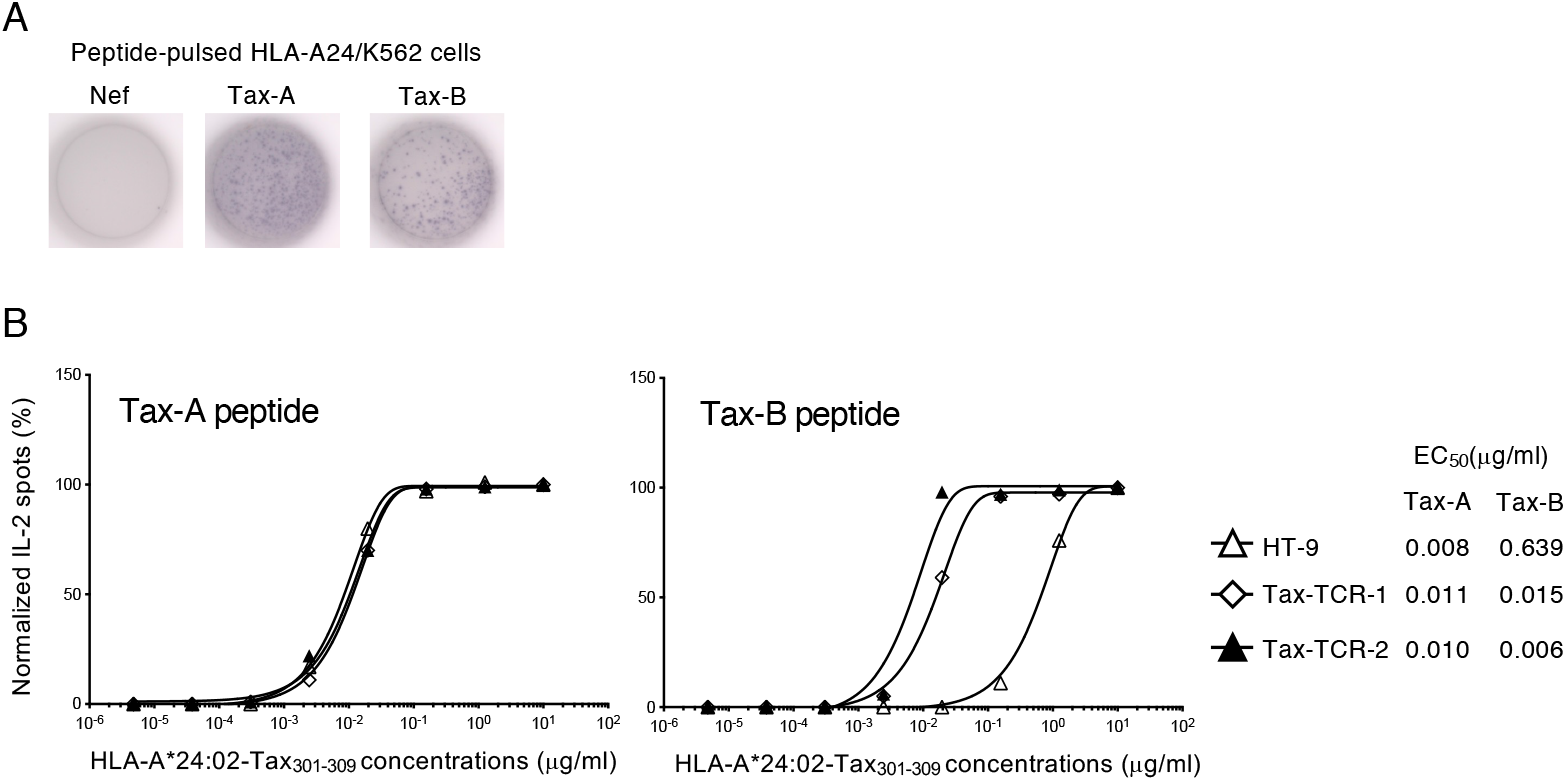
Tax specific TCR (HT-9) shows lower avidity for HLA-A24/Tax-B peptide complex than that of Tax-A. (A) Specific IL-2 production in HT-9-transduced Jurkat cells stimulated with Tax-A and -B peptides. Nef_134-143_ is negative control as HLA-A24-restricted peptide. (B) Comparison of TCR avidity of HT-9 and two newly cloned TCRs for HLA-A24/Tax-A and -B. HLA-A24/K562 cells were pulsed with graded concentrations of the peptides and used to stimulate TCR-transduced Jurkat cells. At least 2 independent experiments were performed.

In conclusion, we demonstrated that mismatches in the viral antigen epitope can lead to inefficient anti-viral T-cell responses. Given the ongoing preclinical and clinical trials targeting HTLV-1 Tax antigen^7,13,19,20^, we would like to recommend checking this viral antigen epitome to ensure optimal immunotherapeutic efficacy.

## Acknowledgements

We are grateful to T. Ueno for TCR-deficient Jurkat T cells and V. Appay for valuable discussion. We also thank M.I. Jahan for critical English editing. This study was supported by grants from the Japan Society for the Promotion of Science (JSPS) KAKENHI (JP20H03724 to Y.S., JP21K08494 to K.S., JP23K14530 to M.T.), Japan Agency for Medical Research and Development (AMED) (JP23fk0410052, JP23wm0325068 and JP23jm0210074 to Y.S.), and JST MIRAI to Y.S. Also, this study is partially supported by grants from the Denka Co. Ltd. To Y.S., and Meiji Seika Pharma Co., Ltd., KM Biologics Co, Ltd. to, K.S. and Y.S. The funders had no role in study design, data collection, data interpretation, or the discussion regarding submission for publication.

## Authorship Contributions

Y.S. contributed to the study conceptualization. K.S. and Y.S. designed research; K.S., M.T. and O.R. performed research; B.T.J.Y. performed bioinformatics analysis; K.S., M.T. and Y.S. analyzed data; M.T., T.S., M.U., Y.Y. and A.U. contributed materials/reagents/analytic tools; K.S. and Y.S. wrote original draft; All authors reviewed and revised the manuscript draft and approved the final version for submission.

## References

1. Bangham CRM. Human T Cell Leukemia Virus Type 1: Persistence and Pathogenesis. Annu Rev Immunol 2018; 36: 43–71.

2. Matsuoka M, Jeang KT. Human T-cell leukaemia virus type 1 (HTLV-1) infectivity and cellular transformation. Nat Rev Cancer 2007; 7(4): 270–80.

3. Kawano N, Shimoda K, Ishikawa F, et al. Adult T-cell leukemia development from a human T-cell leukemia virus type I carrier after a living-donor liver transplantation. Transplantation 2006; 82(6): 840–3.

4. Kawano N, Yoshida S, Kawano S, et al. The clinical impact of human T-lymphotrophic virus type 1 (HTLV-1) infection on the development of adult T-cell leukemia-lymphoma (ATL) or HTLV-1-associated myelopathy (HAM) / atypical HAM after allogeneic hematopoietic stem cell transplantation (allo-HSCT) and renal transplantation. J Clin Exp Hematop 2018; 58(3): 107–21.

5. Harashima N, Kurihara K, Utsunomiya A, et al. Graft-versus-Tax response in adult T-cell leukemia patients after hematopoietic stem cell transplantation. Cancer Res 2004; 64(1): 391–9.

6. Tanaka Y, Nakasone H, Yamazaki R, et al. Long-term persistence of limited HTLV-I Tax-specific cytotoxic T cell clones in a patient with adult T cell leukemia/lymphoma after allogeneic stem cell transplantation. J Clin Immunol 2012; 32(6): 1340–52.

7. Suehiro Y, Hasegawa A, Iino T, et al. Clinical outcomes of a novel therapeutic vaccine with Tax peptide-pulsed dendritic cells for adult T cell leukaemia/lymphoma in a pilot study. Br J Haematol 2015; 169(3): 356–67.

8. Afonso PV, Cassar O, Gessain A. Molecular epidemiology, genetic variability and evolution of HTLV-1 with special emphasis on African genotypes. Retrovirology 2019; 16(1): 39.

9. Jacobson S, Shida H, McFarlin DE, Fauci AS, Koenig S. Circulating CD8+ cytotoxic T lymphocytes specific for HTLV-I pX in patients with HTLV-I associated neurological disease. Nature 1990; 348(6298): 245–8.

10. Kannagi M, Harada S, Maruyama I, et al. Predominant recognition of human T cell leukemia virus type I (HTLV-I) pX gene products by human CD8+ cytotoxic T cells directed against HTLV-I-infected cells. Int Immunol 1991; 3(8): 761–7.

11. Katsuya H, Islam S, Tan BJY, et al. The Nature of the HTLV-1 Provirus in Naturally Infected Individuals Analyzed by the Viral DNA-Capture-Seq Approach. Cell Rep 2019; 29(3): 724–35 e4.

12. Sugata K, Yasunaga J, Mitobe Y, et al. Protective effect of cytotoxic T lymphocytes targeting HTLV-1 bZIP factor. Blood 2015; 126(9): 1095–105.

13. Kawamura K, Tanaka Y, Nakasone H, et al. Development of a Unique T Cell Receptor Gene-Transferred Tax-Redirected T Cell Immunotherapy for Adult T Cell Leukemia. Biol Blood Marrow Transplant 2020; 26(8): 1377–85.

14. Tan BJ, Sugata K, Reda O, et al. HTLV-1 infection promotes excessive T cell activation and transformation into adult T cell leukemia/lymphoma. J Clin Invest 2021; 131(24).

15. Ueno T, Fujiwara M, Tomiyama H, Onodera M, Takiguchi M. Reconstitution of anti-HIV effector functions of primary human CD8 T lymphocytes by transfer of HIV-specific alphabeta TCR genes. Eur J Immunol 2004; 34(12): 3379–88.

16. Furukawa Y, Kubota R, Eiraku N, et al. Human T-cell lymphotropic virus type I (HTLV-I)-related clinical and laboratory findings for HTLV-I-infected blood donors. J Acquir Immune Defic Syndr 2003; 32(3): 328–34.

17. Tanaka Y, Nakasone H, Yamazaki R, et al. Single-cell analysis of T-cell receptor repertoire of HTLV-1 Tax-specific cytotoxic T cells in allogeneic transplant recipients with adult T-cell leukemia/lymphoma. Cancer Res 2010; 70(15): 6181–92.

18. Sanchez-Trincado JL, Gomez-Perosanz M, Reche PA. Fundamentals and Methods for T- and B-Cell Epitope Prediction. J Immunol Res 2017; 2017: 2680160.

19. Tezuka K, Xun R, Tei M, et al. An animal model of adult T-cell leukemia: humanized mice with HTLV-1-specific immunity. Blood 2014; 123(3): 346–55.

20. Suzuki S, Masaki A, Ishida T, et al. Tax is a potential molecular target for immunotherapy of adult T-cell leukemia/lymphoma. Cancer Sci 2012; 103(10): 1764–73.

21. Morita S, Kojima T, Kitamura T. Plat-E: an efficient and stable system for transient packaging of retroviruses. Gene Ther 2000; 7(12): 1063–6.

